# Extreme Small-World, Modular, and Rich-Club Topology of Single-Neuron Networks in Mouse Primary Visual Cortex

**DOI:** 10.1101/2025.08.13.670013

**Authors:** Siqi Tu, Xueting Li, Longsheng Jiang, Guihua Xiao, Jia Liu

## Abstract

Understanding whether the canonical topologies of macroscale connectomes, such as small-world architecture, hub dominance, rich-club cores, and modularity, extend to local cortical microcircuitry has remained challenging due to limitations in simultaneously recording large neuronal populations *in vivo*. Here, using ultra-large-scale, high-resolution calcium imaging, we tracked spontaneous activity from approximately 2,000 neurons across the mouse primary visual cortex (V1). Across multiple mice and correlation thresholds, V1 neuronal networks exhibited hallmark characteristics of efficient brain organization, but in a markedly intensified form compared to macroscopic brain networks. Local clustering coefficients remained an order of magnitude above random levels, while characteristic path lengths approached those observed in random networks, yielding an exceptionally high small-world index that substantially exceeded typical values previously reported at macroscopic scales. Degree distributions followed a power-law, identifying highly connected hub neurons whose interconnections formed a robust rich-club integrative core. Community detection analyses showed robust modularity upon pruning weak connections, indicating functionally specialized neuronal clusters interconnected predominantly through hubs. These findings provide one of the first direct *in vivo* evidence that single cortical microcircuits not only recapitulate but intensify network topologies observed at macroscopic scale, implying evolutionarily conserved design principles underlying brain organization from neurons to systems.

## Introduction

Even in the absence of external stimuli, neurons in the brain exhibit dynamic activity characterized by spontaneous coordinated fluctuations, which collectively reflect intrinsic functional network architectures (Fox & Raichle, 2007; Mitra & Raichle, 2016). Rather than representing random noise or mere epiphenomena, these resting-state activity patterns form structured functional connectivity networks whose topological organization is fundamental to efficient information processing, system stability, and adaptive flexibility (Greicius et al., 2003). Indeed, graph-theoretical analyses consistently reveal canonical architectural motifs in large-scale brain networks, particularly small-worldness, which features prominent local clustering alongside relatively short average path lengths between nodes (Bassett & Bullmore, 2006; Sporns & Zwi, 2004). Such small-world architecture effectively balances segregated specialization and integrated communication, essential for complex brain function. Additionally, brain networks frequently exhibit heavy-tailed (approximately scale-free) degree distributions, characterized by a small number of highly connected hub nodes interspersed among many sparsely connected nodes (Bassett & Bullmore, 2006). Two further prominent organizational motifs are modular community structure, which consists of densely intra-connected groups of nodes (modules) interconnected by sparser inter-module links (Meunier et al., 2010; Sporns & Betzel, 2016), and rich-club organization, wherein a central core of hub nodes is interconnect more densely than would be expected by chance (Ball et al., 2014). Collectively, these topological characteristics are well-documented across numerous macroscopic brain connectomes and are posited to facilitate efficient information transmission, robustness to perturbation, and an optimal balance between localized and global neural processing (Achard et al., 2006; Bullmore & Sporns, 2009; Van Den Heuvel & Sporns, 2011).

However, a fundamental unresolved question is whether these canonical organizational principles observed in macroscale networks are similarly represented at the level of individual neurons within intact cortical microcircuits. To date, most evidence supporting small-world architecture, modular structure, and rich-club organization in neural systems has originated from macroscopic measurements, in which each network node corresponds to an entire brain region or a voxel containing thousands of neurons (Achard et al., 2006; Bullmore & Sporns, 2009; Van Den Heuvel & Sporns, 2011). Such coarse spatial resolution, as typically encountered in resting-state fMRI, inherently obscures single-neuron dynamics. Specifically, the aggregation of neuronal signals may artificially elevate apparent functional connectivity, potentially underestimating the true sparsity of neuronal networks and biasing measured topologies toward more random configurations with lower clustering (Rubinov & Sporns, 2011). Consequently, although human and whole-brain rodent studies have robustly identified modular structure and regional hubs (Bardella et al., 2016; Sporns & Betzel, 2016; Van den Heuvel & Sporns, 2013), they have offered limited insight into whether an *in vivo* local cortical microcircuit intrinsically exhibits similar small-world characteristics. Some supportive evidence has emerged from connectomics and reduced preparations; for instance, the complete neuronal connectome of *C. elegans* (∼279 neurons) exhibits small-world architecture with a densely interconnected rich-club composed of hub neurons (Cook et al., 2019; Towlson et al., 2013), suggesting conservation of certain organizational motifs across mesoscopic and macroscopic scales. Furthermore, developing or cultured neuronal networks exhibit scale-free connectivity and hub neurons that significantly modulate global activity (Downes et al., 2012). Nevertheless, direct evidence demonstrating such complex network topology *in vivo* within mammalian cortical microcircuit has been elusive, primarily due to technological limitations in simultaneously recording neuronal populations at sufficient scale and resolution. Traditional *in vivo* electrophysiological and two-photon imaging methods typically sample only tens to hundreds of neurons simultaneously (Liew et al., 2021), potentially overlooking broader network context and thereby underestimating small-world properties at the single-neuron level. Thus, until recently, it has remained uncertain whether a single cortical microcircuit genuinely demonstrates small-world, modular, and hub-rich organizational motifs characteristic of larger brain networks.

Recent advances in neurotechnology have begun to bridge this scale gap within network neuroscience. Specifically, wide-field calcium imaging, high-density electrode arrays, and other advanced large-scale recording approaches now facilitate the simultaneous monitoring of activity from thousands of neurons distributed across cortical regions *in vivo* (Ren & Komiyama, 2021). Capitalizing on these technological breakthroughs, it is now feasible to reconstruct functional connectivity maps at single-neuron resolution under physiologically relevant conditions. In the present study, we utilize an advanced optical imaging technique, specifically the real-time ultra-large-scale high-resolution (RUSH3D) one-photon calcium imaging system (Fan et al., 2019), to examine the emergent network structure within an intact cortical microcircuit. RUSH3D imaging enables comprehensive capturing of spontaneous neuronal activity across an entire cortical region with both high spatial and temporal resolution, thereby offering an unprecedented mesoscopic perspective into the functional interactions at the level of individual neurons.

In this study, we focused on the mouse primary visual cortex (V1) as a model system, given its well-characterized functional organization and accessibility for wide-field recording (de Vries et al., 2020). V1 is particularly suited for such analyses because neuronal activity recorded during awake resting states, in the absence of sensory stimuli, reliably reflects intrinsic cortical network dynamics (de Vries et al., 2020). Initially, we utilized intrinsic signal optical imaging to localize V1 and subsequently employed RUSH3D imaging (Fan et al., 2019) to precisely record spontaneous neuronal activity from a large neuronal population within this region. This combined approach yielded a comprehensive, high-resolution dataset representing a single-area neuronal network, thus enabling a detailed investigation of its topological properties.

Here we present the first systematic graph-theoretic investigation of cortical resting-state network at single-neuron resolution. Specifically, we reconstructed functional connectivity networks based on spontaneous calcium activity from proximately two thousand V1 neurons and systematically evaluated these networks for canonical topological properties previously described in macroscopic brain networks. We assessed the presence and magnitude of small-world architecture, determined whether the neuronal degree distribution exhibited a heavy-tailed pattern indicative of hub neurons, identified modular community structure, and examined the existence of hub-centered rich-club organization, all at the granularity of single neurons. We hypothesized that even within a local cortical microcircuit, spontaneous neuronal activity would self-organize into networks supporting both local specialization and global integration, reflecting classic small-world properties and related hallmarks of efficient networks organization at the mesoscopic scale. By empirically testing these hypotheses, our study aims to bridge the conceptual and empirical gap between macroscopic connectome principles and the mesoscopic functional architecture of cortical microcircuits. Our findings that neuronal networks within a single cortical area exhibit prominent small-world, modular, and rich-club topologies suggest that fundamental organizational principles of brain networks are conserved across multiple scales, from individual neurons to large-scale brain regions.

## Results

### 1. Resting-state functional connectivity of neuronal network in mouse V1

To investigate the topological characteristics of neuronal networks at the mesoscopic scale, we performed calcium imaging of neuronal activity across the entire visual cortex in mice through a cranial window (Figure 1a). Figure 1a (middle) shows a retinotopic layout of the visual cortex derived from intrinsic signal imaging (ISI) (Siegle et al., 2021), with different colors representing cortical response amplitudes and phases elicited by varying temporo-spatial frequencies, thus delineating the visuotopic structure of cortical areas. Cortical neurons were labeled with a fluorescent calcium indicator, and spontaneous neuronal activity was recorded using the RUSH3D imaging system, enabling wide-field acquisition at single-neuron resolution. Figure 1a (right) illustrates a representative fluorescence image of the cortical surface at single-neuron resolution, capturing neuronal somata within V1. By integrating functional localization via ISI with large-scale calcium imaging, we precisely identified neuronal populations in V1 and their spatial coordinates, facilitating the construction of resting-state functional connectivity networks.

**Figure 1.**
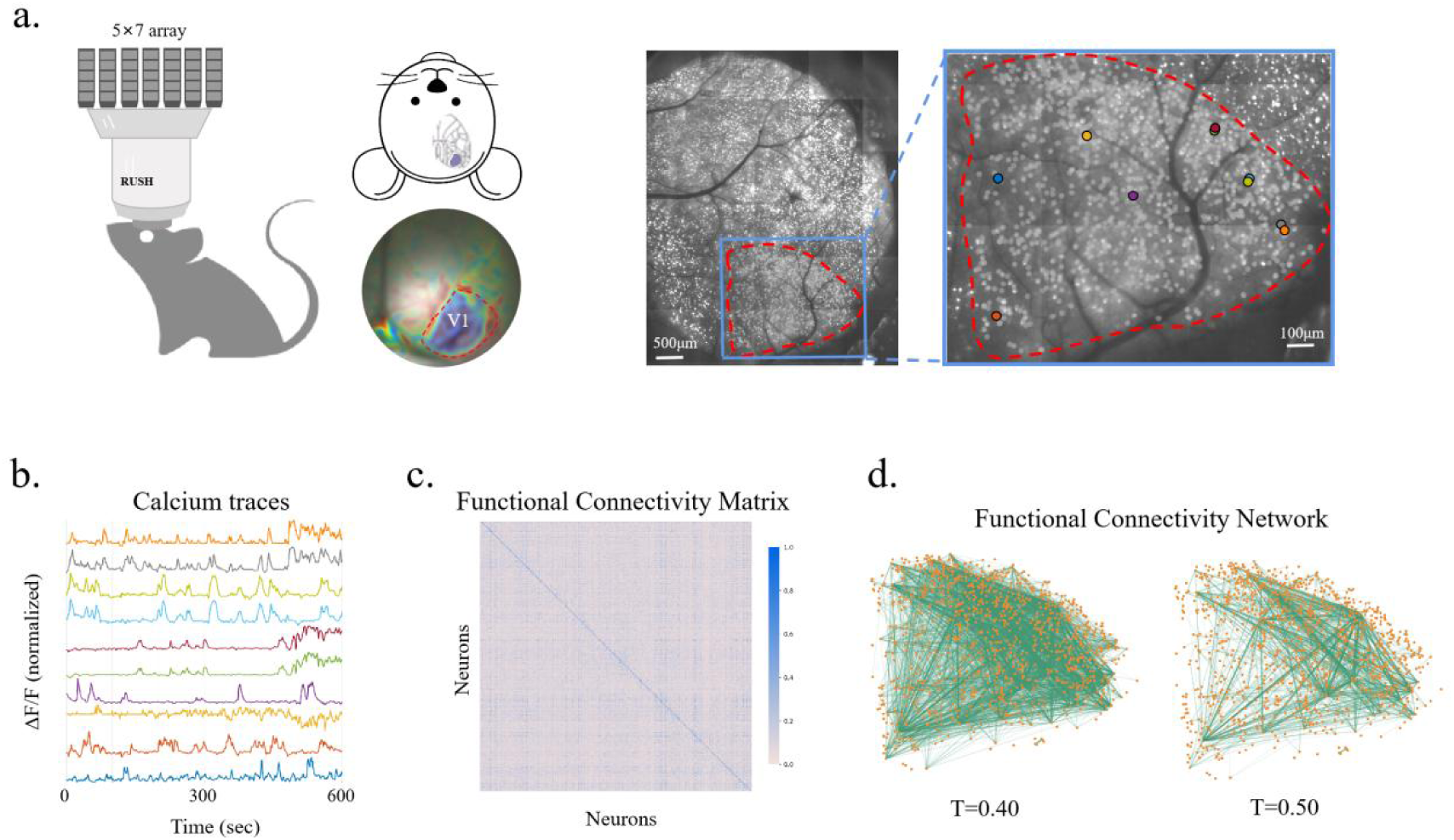
Resting-state functional connectivity of neuronal networks in mouse V1. (a) Left: Schematic illustration of the calcium imaging setup utilized for recording spontaneous neuronal activity in the mouse visual cortex *in vivo*. Middle: Cranial window placement (top) and the ISI map with V1 delineated by the red dashed line (bottom). Colors indicate cortical responses to distinct temporo-spatial frequencies. Right: Representative wide-field fluorescence image of neuronal somata within V1 acquired via calcium imaging; scale bar, 100 μm. (b) Example calcium fluorescence traces recorded from individual neurons in mouse V1. (c) Functional connectivity matrix generated from pairwise Pearson correlation coefficients calculated from spontaneous calcium fluorescence traces. (d) Visualization of functional connectivity networks at two representative correlation thresholds (T = 0.40 and 0.50, respectively). Orange dots represent individual neurons, and green lines denote functional connections.

With the neuronal population within V1 identified, we next extracted spontaneous calcium signals from these neurons during the resting state, in the absence of sensory stimulation. Figure 1b illustrates representative calcium fluorescence traces from a subset of V1 neurons recorded in one mouse over several minutes. Each trace corresponds to the ΔF/F signal from an individual neuron, reflecting fluctuations in intracellular calcium concentrations indicative of neuronal spiking activity. These spontaneous fluorescence signals exhibited intricate temporal dynamics, with neurons firing irregularly and, occasionally, subsets of neurons displaying coordinated bursts or co-fluctuations, indicative of synchronous network activity during the resting state. To quantitatively characterize functional r interactions among neurons, we computed the pairwise Pearson correlation coefficients between calcium time series for all neuronal pairs (Figure 1c). Consequently, each neuron represented a node in the constructed functional connectivity network, and an edge between two nodes was defined when the correlation between two neurons exceeded a predetermined threshold. Notably, given the large-scale recordings encompassed an average of nearly two thousand neurons simultaneously (8 mice tested; mean number of neurons:1898; range: 824-2615; SD = 664.2), the network provided a comprehensive and unbiased representation of spontaneous neuronal interactions within V1 during resting state.

Figure 1d depicts representative examples of neuronal networks constructed at different correlation thresholds (T), reflecting varying degrees of sparsity. At a relatively permissive threshold (T = 0.40, Figure 1d, left), the resultant network exhibited dense interconnectivity. As the correlation threshold was elevated to 0.50, weaker functional connections were progressively eliminated, leading to a sparer connectivity profile. Crucially, despite significant pruning of weaker edges, core network structures remained discernible; clusters of neurons persisted as interconnected modules, and hub-like neurons retained numerous robust connections. This topological organization was consistently observed across all mice teste (Supplemental Figure 1), suggesting a stable functional architecture underlying spontaneousV1 activity and highlighting the existence of fundamental modular subnetworks within the cortical microcircuit. To further characterize these structural features, we next quantitatively evaluated canonical network metrics, specifically emphasizing the balance between high local clustering and short path lengths in the V1 network.

### 2. Small-World architecture characterized by high clustering and short path lengths

We quantified two fundamental network properties in the V1 network: the clustering coefficient (C), which reflects the degree of local interconnectedness, and the characteristic path length (L), which measures the average shortest path between pairs of nodes. Subsequently, we compared these metrics with those obtained from degree-matched random networks. Detailed small-world network properties across the group of eight mice are presented Figure 2a, while Figure 2b s summarizing the small-world indices computed for each individual mouse. These analyses illustrate the consistency of small-world architecture across mice.

**Figure 2.**
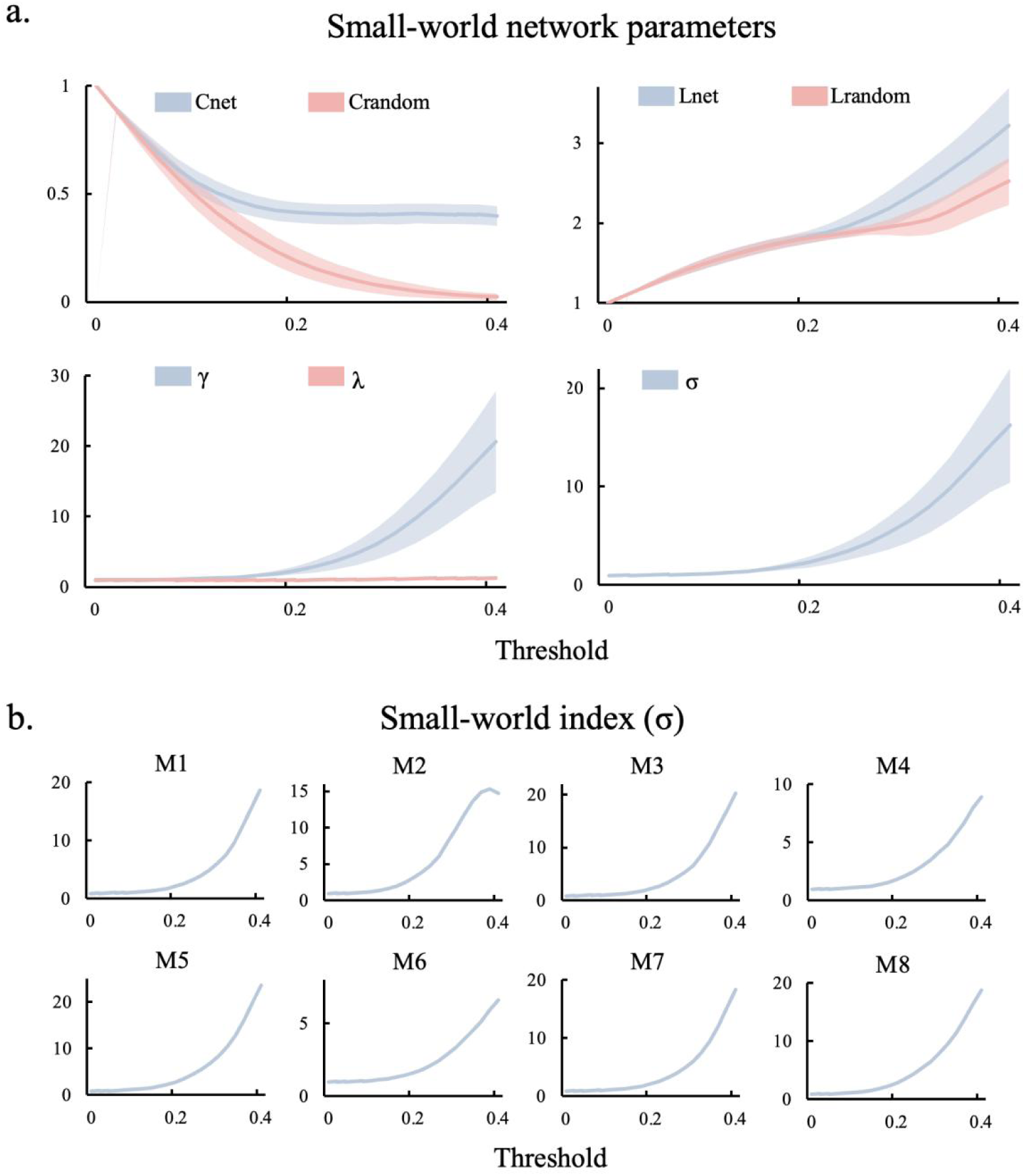
Small-world architecture. (a) A quantitative analysis of small-world network properties was conducted across varying correlation thresholds. The upper-left panel shows the clustering coefficient (C) for neuronal networks (*C_net_*, blue) in comparison to the matched random networks (*C_random_*, pink). The upper-right panel displays the characteristic path length (L) for neuronal networks (*L_net_*, blue) relative to the random networks (*L_random_*, pink). The lower-left panel illustrates the ratios of clustering coefficients (*γ =C_net_/C_random_*) and path lengths (*λ =L_net_/L_random_*). The pronounced increase in clustering, alongside relatively stable path lengths, indicates that the small-world architecture in V1 was primarily driven by enhanced local clustering. The lower-right panel presents the small-world index (*σ*), which consistently exceeded 1 across all thresholds above 0.02, peaking approximately 18 at T = 0.40, further confirming the robust small-world architecture. Shaded areas represented ±1 standard deviation (SD) across the eight mice. (b) Small-world indices (*σ*) computed for individual mice are shown, illustrating the consistency and robustness of the small-world architecture across mice.

Classic graph theory characterizes small-world architecture as the coexistence of exceptionally high clustering with path lengths that are comparable to those observed in random networks. To assess this, we computed the normalized clustering coefficient *γ =C_net_*/*C_rand_* and normalized path length *λ =L_net_*/*L_rand_*, where the subscript “net” refers to the V1 network and “rand” refers to the corresponding values from degree-matched random networks that preserve both node degrees and edge counts (for details, see Methods). The small-world index (σ) was then computed as the ratio of the normalized clustering coefficient to the normalized path length (*σ* = *γ*/*λ*), which provides a measure of the extent to which a network’s clustering exceeds that of a random networks, once differences in path length are accounted for (Humphries et al., 2006). A small-world index (σ) around 1 or slightly above is typically observed many complex brain networks, whereas substantially higher values indicate a notably more pronounced small-world architecture. To investigate small-world properties at varying network densities, we constructed binary functional networks across a broad range of correlation thresholds (T = 0.02 to 0.40, in increments of 0.02). Figure 2 provides a summary of how clustering and path length, as well as their normalized counterparts, vary as the network density decreases.

The clustering coefficient of the V1 network (*C_net_*, blue) remained consistently high across a broad range of correlation thresholds, in contrast to the clustering observed in the degree-matched random networks (*C_random_*, pink) (Figure 2, upper left). Even as weaker connections were progressively removed at higher thresholds, *C_net_* remained substantial (ranging from approximately 0.20 to 0.40 in absolute terms, SD = 0.046), showing only minimal decline. In contrast, *C_random_* decreased sharply with increasing threshold, as random networks of equivalent sparsity lack extensive clustered connections. For all thresholds above approximately T = 0.02, the observed clustering in the V1 network significantly exceeded that of random networks (paired t-tests, *p* < 0.05, Bonferroni-corrected), indicating that V1 neurons formed highly interconnected local neighborhoods, maintaining high clustering even under sparse connectivity conditions. This is a hallmark of small-world architecture.

The characteristic path length of the V1 network (*L_net_*, blue) increased gradually as the correlation threshold was raised, as the pruning of weaker connections disrupted some of the shortest paths, thereby modestly lengthening the communication routes between neurons (Figure 2a, upper right). Over the threshold range from 0.2 to 0.4, *L_net_* increased from approximately 1.8 to 3.2, representing the average number of steps between neuron pairs. In contrast, the characteristic path length of the degree-matched random networks (*L_random_*, pink) exhibited expected behavior for random networks, showing a slight increase as edges were removed. Notably, at lower and moderate thresholds (T < 0.3), *L_net_* remained comparable to *L_random_*, indicating that the V1 network maintained integration efficiency similar to that of random network with equivalent density. At higher threshold levels (T ≥ 0.30), *L_net_* became slightly, but significantly, larger than *L_random_* (*p* < 0.05, Bonferroni-corrected), reflecting a modest reduction in communication efficiency in the V1 network when only the strongest connections remained. Crucially, even at the highest sparsity levels, the path length of the V1 network remained on the same order of magnitude as that of random networks (within a factor of approximately 1.1–1.4 of *L_random_*), indicating V1 network preserved short path lengths between neurons despite substantial pruning of weak connections.

Figure 2a (lower left) presents the ratio metrics (*γ* and λ) to explicitly illustrate the small-world architecture of the V1 network. The normalized clustering coefficient (*γ*, blue) remained exceedingly high, approximately 5 at lower thresholds and increasing sharply to nearly 20 at the highest thresholds, indicating that the local clustering in the V1 network was orders of magnitude greater than the random network expectation. In contrast, the normalized path length (λ, pink) remained very close to 1 across all thresholds, with only a slight increase at the highest threshold, indicating that the global path length in the V1 network was similar to that of random networks. That is, as the network was pruned to retain only the strongest connections, the relative reduction in clustering observed in random networks was substantially greater than in the V1 network, while the corresponding increase in path length was minimal. This divergence between *γ* and λ highlights the V1 network’s ability to maintain highly efficient local clustering while preserving overall integration without significant compromise.

Figure 2a (lower right) shows the small-world index (σ), which consolidates the aforementioned effects into a single measure. We observed that σ remained consistently above 1.0 across all thresholds tested, confirming the robust presence of small-world architecture at various levels of sparsity. At lower thresholds, σ was elevated (approximately 5–10) and increased substantially as threshold was raised, peaking around 20 at T = 0.40. Practically, this suggests that the V1 network demonstrated a nearly twentyfold greater small-world effect compared to random networks at equivalent sparsity. Notably, this pronounced small-world architecture was consistently observed across individual mice (Figure 2b), highlighting the biological reproducibility and robustness of the network’s organization.

The small-world index (∼20) in V1 indicated a network with prominent local cliques (*γ* ≫ *1*) and efficient global integration (λ ≈ 1), reflecting an exceptionally strong small-world effect within this mesoscopic neuronal network. For comparison, previous studies of whole-brain functional networks typically reported σ ranging from 2 to 5, driven by moderately elevated clustering and nearly random-like path lengths (*γ* > 1, λ ≈ 1) (Bassett & Bullmore, 2006; van den Heuvel et al., 2008; Watts & Strogatz, 1998). Thus, the mesoscopic V1 network achieved an order of magnitude stronger small-world effect than macroscopic brain networks, effectively balancing tight local clusters and efficient long-range connectivity. This efficiency likely depends on hub neurons, which acting as integrative shortcuts between distinct neuronal clusters, combined with a modular structure that promotes high within-module clustering. To further explore the origins of the V1 network’s extreme small-world architecture, we next examined whether this organization arises from a hub-and-module organization.

### 3. Hub neurons and scale-free degree organization

A “hub” is typically defined as a node hat exhibits a significantly higher number of connections compared to the average node in the network, functioning as a central point for information flow (Albert & Barabási, 2002). Networks characterized by heavy-tailed or scale-free degree distributions (*P* (*k*) = *k*^−α^) often feature such hubs, where the majority of nodes possess only a few connections, while a small subset of nodes, the hubs, are highly connected. Indeed, in the V1 network, the degree distribution followed an approximate power-law, particularly at higher correlation thresholds. Figure 3a presents the node degree distributions on log–log axes for two representative sparsity levels (T = 0.35 and T = 0.40). In both cases, the distributions exhibit a strikingly linear trend, consistent with the hallmark pattern of scale-free organization (Bassett & Bullmore, 2006). The fitted power-law exponents (α ≈ 1.04 to 1.32) reflected a distinctly heavy-tailed degree distribution, in which a small subset of neurons exhibited exceptionally high connectivity, while the vast majority maintained relatively few connections. This pattern represents a pronounced departure from the Poisson-like degree profile characteristic of random networks, underscoring the presence of hub neurons within the V1 network.

**Figure 3.**
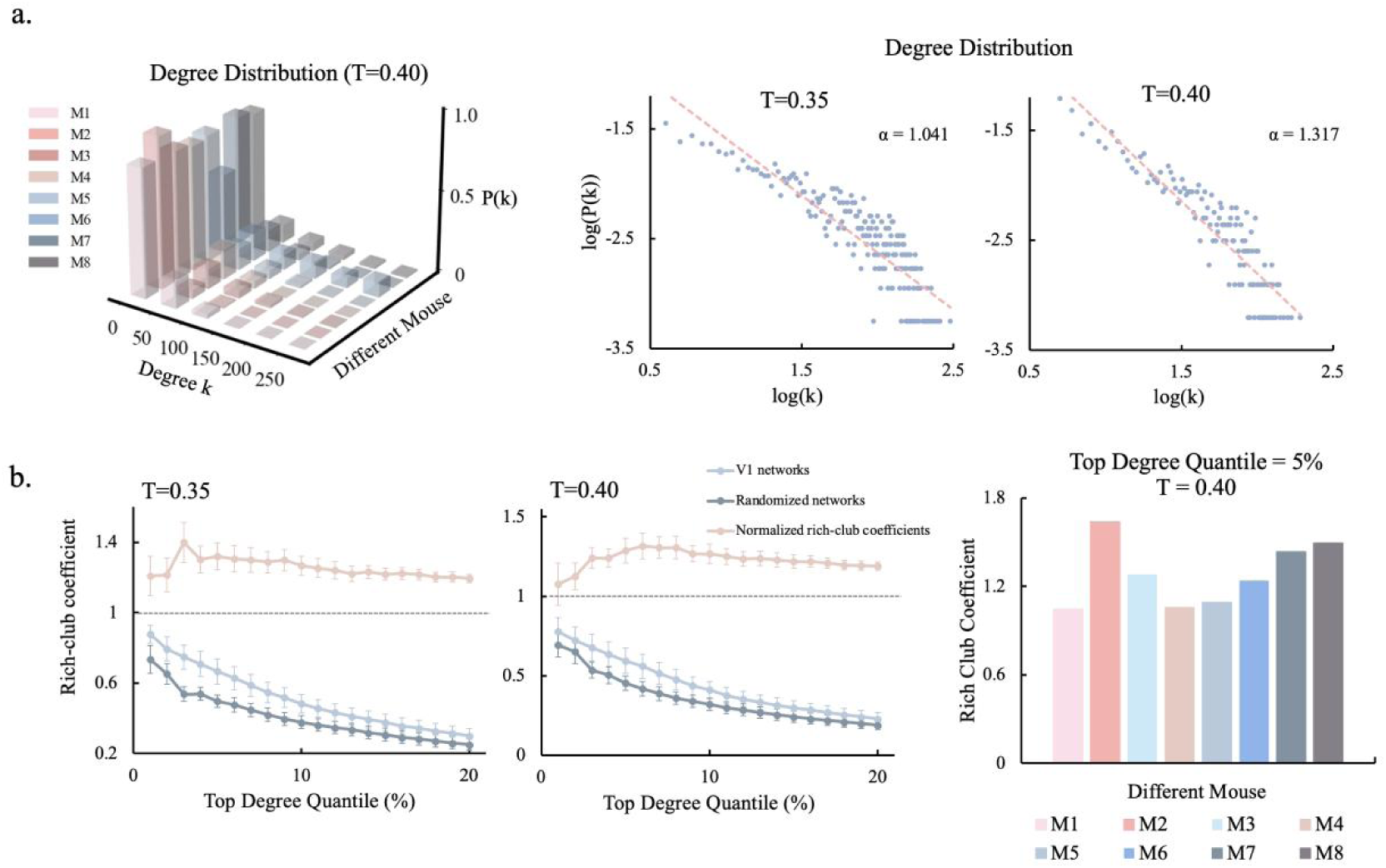
Hub neurons and scale-free organization. (a) Degree distributions of neuronal networks at varying correlation thresholds across different mice. The left panel shows degree distributions at T = 0.4 for eight individual mice, with each mouse represented by a distinct color, illustrating consistent heavy-tailed, scale-free patterns. The upper-right panel displays log-log plots of node-degree (k) distributions at correlation thresholds T = 0.35 and 0.40, further emphasizing the scale-free characteristics. Each distribution fits a power-law, with the fitted exponents (*α*) indicated by the red dashed lines. (b) Rich-club organization. The lower-left panel illustrates rich-club coefficients (*ф*) at thresholds T = 0.35 and T = 0.40. Light blue: V1 networks; dark grey: corresponding randomized networks; pink: normalized rich-club coefficients (*ф_net_*(*k*)*ф_rand_*(*k*)). Notably, normalized coefficients consistently exceeded 1 for the top-degree quantiles (3% – 20%). Data points represent the mean values ±1 standard error (SE). The right panel presents normalized rich-club coefficients for eight individual mice at T = 0.40, focusing on the top 5% quantile.

The emergence of scale-free degree distributions within the V1 network suggests a hub-centric organizational structure, where certain neurons (i.e., hubs) exhibit strong co-activation with a substantial portion of the network, facilitating short communication distances and effective integration (Eguiluz et al., 2005). Figure 3b further demonstrates that these hub neurons preferentially interconnect, forming a rich-club organization that effectively establishes a high-capacity backbone for information flow within V1 (Bullmore & Sporns, 2012; Van Den Heuvel & Sporns, 2011). The rich-club organization is quantified by plotting the rich-club coefficient (*ф(k)*) as a function of degree rank for V1 networks, compared to random networks. ф(*k*) measures connectivity density among nodes with a degree exceeding k. As shown in Figure 3b, *ф_net_*(*k*) for V1 networks consistently surpassed *ф_rand_*(*k*) for random networks across a broad range of high-degree percentiles (approximately the top 3%–20%). This rich-club organization is further supported by the statistical analyses, where the normalized rich-club coefficients consistently exceeded 1 (*p* < 0.01, Bonferroni-corrected).

Previous studies of macroscopic brain networks have similarly reported the presence of rich-club organization, with highly interconnected hubs forming a central integrative backbone (Ball et al., 2014). Our findings extend this concept to local cortical microcircuits, indicating that neurons with highly connectivity in V1 preferentially form links with one another, thereby reinforcing their roles in network integration. This scale-free, hub-rich organization may account for the observed short characteristic path lengths, even as network sparsity increases, while simultaneously maintaining high clustering, thus contributing to the exceptionally large small-world index (σ) observed. To further investigate the network’s structure, we next examined its community organization to determine whether neurons were arranged into distinct functional modules.

### 4. Modular community structure

In complex brain networks, modularity refers to the organization of nodes into groups that are densely interconnected within each group, while exhibiting sparser connections between groups. Within each module, neurons form tightly integrated circuits specialized for specific functions (high segregation), whereas sparse inter-module connections facilitate efficient global communication (integration). Considering the prominent small-world and hub-rich organization observed, we hypothesized that these network properties were underpinned by a modular community structure. To test this, we conducted modularity analyses to investigate whether the high clustering observed in V1 was indicative of distinct functional modules.

Figure 4a shows the full weighted functional connectivity matrix for all recorded V1 neurons (N = 1756) from a representative mouse, where each entry represents the correlation coefficient between neuron pairs. Visual inspection suggested the presence of an underlying modular structure, with distinct patches of high correlation (dark regions) along the diagonal, indicating subsets of neurons that are more strongly interconnected within their respective groups. To quantify these communities, we applied a correlation thresholding (T = 0.4) and performed community detection. Figure 4b displays the binary adjacency matrix, sorted by community assignment, which clearly displays distinct blocks of intra-connected neurons along the diagonal, providing further evidence of a modular network structure.

**Figure 4.**
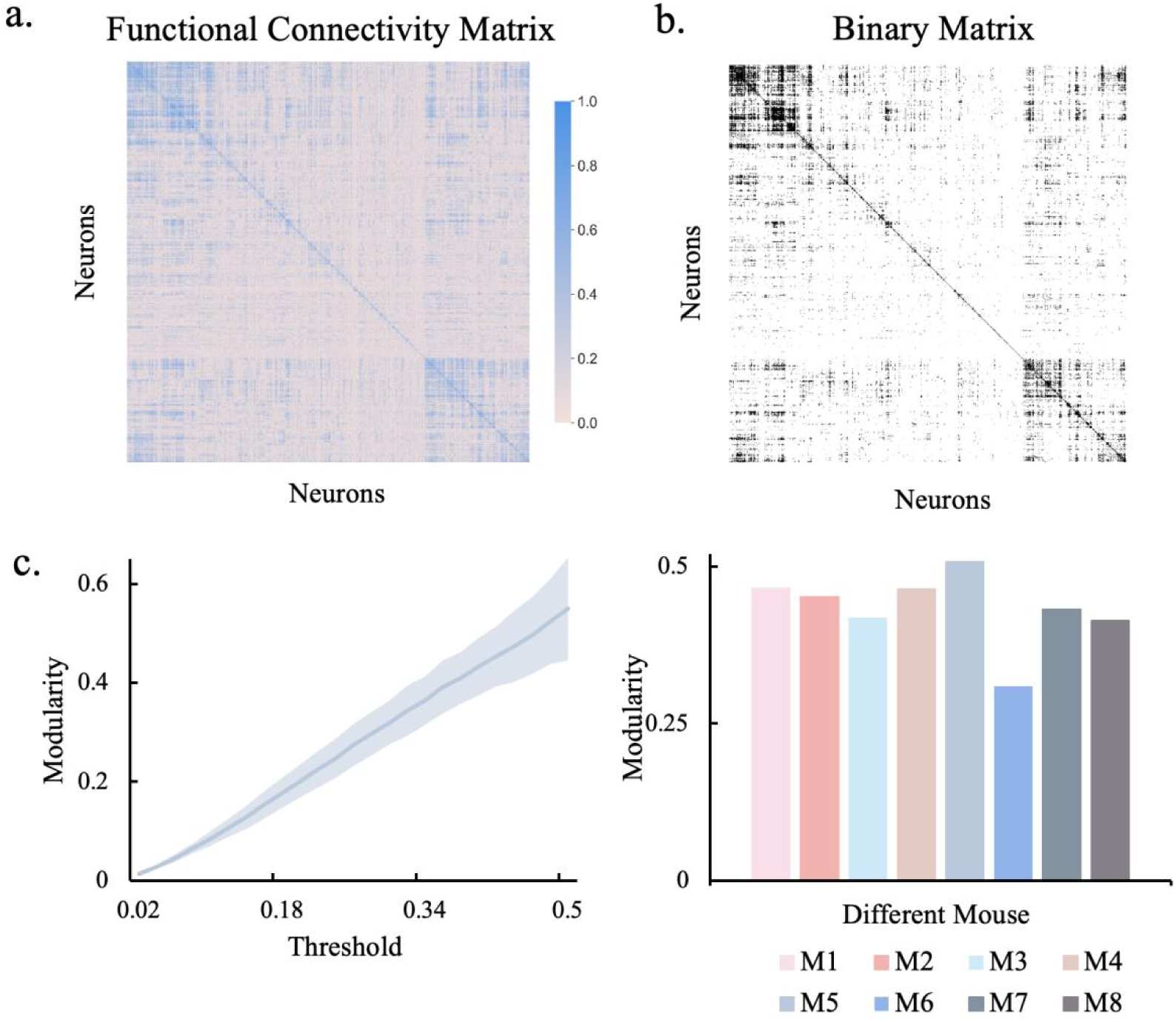
Modularity of mouse V1 networks. (a) Weighted functional connectivity matrix (N = 1756 neurons) illustrating the correlation strengths between neuronal pairs. Darker colors indicate stronger correlations. (b) Binary connectivity matrix at a correlation threshold of T = 0.4, revealing distinct modular community structures. Neurons are sorted according to modularity detection, with dense intra-module connectivity represented by black blocks along the diagonal and sparse inter-module connectivity indicated by off-diagonal white areas. (c) Modularity index (Q) as a function of correlation thresholds (left) and across individual mice at T = 0.4 (right). Modularity increased significantly at higher thresholds, reflecting a more robust community structure. The blue line represents the mean Q ±1 SD across eight mice. All eight individual mice consistently exhibited high modularity, confirming the robustness of the modular structure across mice.

Figure 4c presents the modularity quality index (Q, Newman’s modularity) as a function of correlation thresholds. Modularity Q quantifies the strength of community structure, with higher values indicating more distinct modules. We observed that Q increased significantly with higher thresholds, as weaker connections were progressively removed. Notably, for correlation thresholds greater than approximately T ≥ 0.36, modularity consistently exceeded Q > 0.3 on average, indicating statistically significant community structure (*p* < 0.05, Bonferroni-corrected). This suggests that as weaker connections were pruned, the network became more distinctly modular. To assess consistency across mice, we evaluated modularity at a fixed threshold (T = 0.4) for eight individual mice. As shown in the lower panel of Figure 4c, all mice exhibited high modularity, despite individual variability, further confirming the robustness of the modular structure.

The emergence of densely interconnected modules with sparse inter-module connections suggests that the V1 network is organized into specialized neuronal clusters. Neurons within each module likely share common inputs or functional roles, resulting in strong intra-module correlations. The sparse inter-module connections indicate that communication between modules occurs via critical integrative nodes, such as the hub neurons identified earlier. This structure reflects fundamental principles of efficient brain network architecture: segregated specialization within modules, coupled with selective integrative connections between them.

In summary, the modular structure suggests that V1 neurons form semi-independent functional units specialized for local processing, embedded within a network framework that is mediated by hub-driven integrative connections. This synergistic organization, combining clustered communities, integrative hubs, and a rich-club core, defines V1 as a highly efficient neuronal network that optimally balances local specialization and global integration.

## Discussion

This study investigated whether hallmark topological features previously identified in macroscopic brain networks similarly manifest at the mesoscopic scale within intact cortical microcircuits. Our findings demonstrated that spontaneous functional connectivity among approximately 2,000 neurons in mouse V1 exhibits a pronounced small-world architecture, characterized by exceptionally high local clustering coefficients and relatively short path lengths. Furthermore, these neuronal networks display heavy-tailed degree distributions indicative of prominent hub neurons, forming robust rich-club cores, along with distinctly community structures. Thus, the V1 neuronal networks recapitulate canonical network properties extensively documented at macroscopic scales, but in considerably more extreme manner. By directly validating these network attributes at single-neuron resolution *in vivo*, our results bridge the conceptual and empirical gap between macroscopic connectome principles and the mesoscopic functional architecture of cortical microcircuits. Consequently, this study provides providing direct empirical evidence that conserved organizational principles govern brain network architecture across multiple spatial scales, extending from individual neurons to macroscopic brain regions, thereby enabling an optimal balance between functional segregated specialization and integrated communication.

Our findings are consistent with previous studies that have firmly established small-world architecture, modular structure, and rich-club organization in human and rodent brain networks at the macroscopic scale (Achard & Bullmore, 2007; Bardella et al., 2016; Sporns & Betzel, 2016; Van Den Heuvel & Sporns, 2011). However, these studies typically employed coarse spatial resolutions in which each network node represents an entire brain region or voxel comprising thousands of neurons (Eguiluz et al., 2005; Liska et al., 2015). Although analyses at the mesoscopic scale in cultured neuronal networks (Downes et al., 2012), model organisms (Towlson et al., 2013), and sparse *in vivo* electrophysiological recordings (Nigam et al., 2016) have suggested analogous network attributes, such evidence has largely been indirect. In contrast, the present study directly demonstrates, through high-resolution, single-neuron network analysis, that neurons within a single cortical region spontaneously organize into complex networks following the same canonical organizational principles. The observed self-similarity of brain organization implies hierarchical repetition of efficient network motifs, suggesting that evolutionary and developmental constraints may have driven the emergence of optimal network patterns from microcircuits up to entire brain systems (Bullmore & Sporns, 2012; Kaiser & Varier, 2011).

A particularly notable finding of our study is the exceptionally high small-world index (σ*∼*20) observed in the V1 neuronal network, which markedly surpasses values commonly reported for brain networks at macroscopic scale (Achard & Bullmore, 2007; Bassett & Bullmore, 2006). Such pronounced small-world architecture likely emerges due to neurons forming dense local interconnections that preserve high clustering, complemented by strategically positioned long-range connections that maintain short communication paths. This network configuration achieves an optimal balance between functional segregation, supported by locally clustered modules specialized for distinct computational tasks, and global integration, enabling efficient information transmission across the network. This balance may be realized through an economical wiring strategy, wherein dense local clustering predominantly maintained via abundant short-range connections between adjacent neurons. Such short-range connectivity is metabolically “cheap” and reflective of the densely recurrent organization intrinsic to cortical microcircuits. Conversely, a limited number of long-range connections effectively sustain global integration. This economical organization aligns with theoretical principle proposing that brain networks strive to maximize communication efficiency while minimizing wiring cost (Bullmore & Sporns, 2012). The present study provides empirical support for this principle by demonstrating that cortical networks at the single-area, single-neuron resolution integrate dense local clustering with strategically positioned long-range shortcuts, thereby achieving exceptionally high small-world efficiency.

In addition to the small-world architecture, we observed that the neuronal connectivity distribution followed a scale-free pattern, characterized by a small subset of neurons (hubs) exhibiting significantly higher connectivity. This hierarchical structure, with a few highly connected nodes positioned above a large base of sparsely connected ones, is recognized for generating complex dynamics and promoting resilience in networks, which is a hallmark of efficient communication networks (Albert et al., 2000). Our findings therefore extend the principles of hub-rich organization, commonly observed at macroscopic scales (Eguiluz et al., 2005) by demonstrating that this hub-rich organization is an inherent feature of the neuronal network as well. Furthermore, this *in vivo* evidence of hub neurons corroborates previous *in vitro* studies of cortical networks and developmental research, which suggest that certain hub neurons play a crucial role in driving network-wide events or synchrony (Downes et al., 2012; Nigam et al., 2016). Our results further extend these findings by illustrating the prevalence and functional significance of hub neurons within an awake resting-state cortex. Although speculative, specific neuron types, such as large pyramidal cells with extensive axonal arborizations that link multiple local targets or inhibitory interneurons, might inherently function as hubs connecting distinct clusters. Further studies should explore the cell-type identity of these hubs.

Crucially, hub neurons preferentially form connections with one another, creating a rich-club organization, which has been identified in macroscopic brain networks as a core subnetwork responsible for ensuring robust communications (Collin et al., 2014; Van Den Heuvel & Sporns, 2011). From a systematic perspective, the emergence of rich-club organization at the mesoscopic scale indicates that even within V1, information processing likely depends on this high-capacity backbone to synchronize neuronal ensembles during active cortical states or demanding cognitive processes (see also (Nigam et al., 2016). For instance, during states of alertness or sensory processing, these hub neurons may quickly propagate modulatory signals or coordinate the recruitment of multiple ensembles. Additionally, the rich-club organization accounts for the network’s pronounced small-worldness, as, even after substantial pruning, interconnected hubs sustain integration across otherwise segregated modules, thus preserving short path lengths and high clustering.

Another important network characteristic is modularity within the V1 network, which suggest that neurons in V1 do not form a single, homogeneous network; instead, they are organized into multiple distinct modules. This modular organization mirrors that observed at the macroscopic scale (Sporns & Betzel, 2016), indicating that brain networks are hierarchically modular, with smaller neuronal groups nested within larger modules (e.g., cortical columns or areas), which in turn are incorporated into broader whole-brain networks. In the V1 network, this modular structure likely emerges from stable, functionally similar neuronal groups that process related features or belong to the same canonical microcolumn or layer-specific ensemble, as evidenced by local clusters of co-active neurons in the sensory cortex (Cossell et al., 2015). Our study further reveals that the modular structure becomes more prominent only after pruning weak correlations, implying that robust functional connections establish the core of the modules, while weak connections may blur module boundaries by connecting nodes indiscriminately. This threshold-dependent emergence of modularity implies that strong correlations are crucial for the stability of modules, whereas weaker correlations contribute to the network’s dynamic flexibility. Future studies comparing resting and task-driven states will provide further insights into the dynamics of modularity and functional specialization.

Despite its efficiency, the architecture of the V1 network also implies inherent vulnerabilities. Dysfunction or loss of a few hub neurons could profoundly disrupt global integration, mirroring observations at the macroscopic scale in neurological disorders such as Alzheimer’s disease, schizophrenia, and epilepsy (Van den Heuvel & Sporns, 2013). Specifically, deficits at the mesoscopic level, such as excessive pruning of hub neurons during development, selective degeneration of hub neurons, or seizure-induced overload of hub neurons, could similarly destabilize cortical circuits. Consequently, interventions aimed at protecting or modulating these critical neurons may offer substantial benefits for preserving network integrity. Future studies should test these predictions by selectively silencing or ablation of hub neurons using neuron-type-specific lesion or silencing techniques, such as optogenetic photoinhibition, chemogenetic inactivation, or calcium-dependent ablation, and by assessing both network integrity and associated behavioral outcomes.

A fundamental unanswered question is how these network attributes self-organize during development. Theoretical studies propose that mechanisms such as preferential attachment, spatial constraints, Hebbian plasticity, and activity-dependent pruning sculpt initial random connectivity into structured networks exhibiting small-world, modular, and rich-club properties (Barabási & Albert, 1999; Sur & Leamey, 2001; Zamora-López et al., 2016). Computational models operating near criticality (Liu et al., 2025) replicate these integrated features, suggesting that the V1 network reflects an energy-efficient optimization between local specialization and global integration (Artime et al., 2024; Bullmore & Sporns, 2012). Moreover, evolution likely reinforces this design, and intrinsic cellular factors, such as extensive dendritic arbors, heightened excitability, or specific gene expression profiles, may predispose certain neurons to function as hubs (Van den Heuvel & Sporns, 2013). Future developmental and computational models should seek to replicate the network motifs identified in this study, offering deeper insight into the principles governing cortical self-organization.

In conclusion, our *in vivo* investigation of mouse V1 network reveals that a cortical microcircuit at single-neuron resolution exhibits canonical brain network architecture, such as pronounced small-worldness, robust modularity, and a prominent rich-club core, in a more intensified form compared to macroscopic networks. This structural homology across scales provides support for universal design principles underlying brain network efficiency, thereby encouraging future multi-scale computational models that incorporate realistic neuron-level graphs within each brain region. Such models will facilitate the exploration of how local oscillations, avalanches, or lesions propagate through larger connectomes. Additionally, developmental studies involving longitudinal imaging of young animals could help elucidate whether small-world, modular, and rich-club properties are genetically pre-specified or sculpted by experience. Finally, since hub and rich-club disruptions are hallmark features of brain disorders at the macroscopic scale, detecting analogous mesoscopic alterations in disease models could inform therapeutic strategies aimed at preserving or restoring microcircuit clustering and hub integrity.

## Methods

### Animal preparation

All experimental procedures involving live animals were conducted in strict accordance with Tsinghua University guidelines and approved by the Institutional Animal Care and Use Committee (IACUC) of Tsinghua University, Beijing, China. Mice were housed in the Laboratory Animal Research Center at Tsinghua University under standardized conditions, including a temperature of 24°C, 50% relative humidity, and a reversed light/dark cycle (lights on from 7:00 AM to 7:00 PM).

Mice used in this study were generated by crossing Rasgrf2-2A-dCre (JAX 022864) with Ai148 (TIT2L-GC6f-ICL-tTA2)-D (JAX 030328), resulting in Cre-dependent expression of GCaMP6f (Jackson Laboratory). Adult male and female mice, aged 8–24 weeks and weighing 20–30 g, were housed in groups in standard plastic cages with bedding under controlled environmental conditions. Experimental mice were administered intraperitoneal injections of trimethoprim (TMP, 0.25 mg/g, Sigma) for two consecutive days. Prior to surgery, mice were provided ad libitum access to food and water. Post-surgery, they were housed individually and habituated to the head-fixed system for at least one week before recording.

### Craniotomy operation for wide-field imaging

Mice were anesthetized with isoflurane (4% for induction and 1–2% for maintenance, delivered at 1 mL/min O₂) using an anesthesia machine (R520 IP, RWD Life Science Co.) and meloxicam (5 mg/kg, subcutaneously) was administered for analgesia. Surgical instruments and workstations were sterilized through high-temperature, high-pressure autoclaving before each procedure. Mice were carefully positioned and secured in a stereotaxic apparatus (68018, RWD Life Science Co.), and body temperature was maintained at 37.0 °C using a heating pad (RWD Life Science Co.).

To prepare the mice for all-optical experiments, the scalp was excised, and a 7-mm-diameter craniotomy was performed over the right monocular visual cortex (centered at 1.3 mm lateral and 2.5 mm anterior to lambda), which processes visual information from the left eye. A chronic cranial imaging window was implanted to replace the excised skull and was secured using Krazy Glue (Elmer’s Products Inc.). An aluminum headplate with a 10-mm-diameter circular imaging well was then affixed to the skull over the cranial window using dental cement. This setup enabled wide-field imaging of single-neuron activity across the visual cortex.

Postoperatively, mice received subcutaneous injections of the anti-inflammatory drug flunixin meglumine (1.25 mg/kg, Sichuan Dingjian Animal Medicine Co.) for five consecutive days. They were then allowed to recover for at minimum of seven days, during which the integrity of the cranial window and the quality of calcium signals were evaluated. Only mice that fully recovered and showed no signs of distress were included in subsequent experiments. Mice were observed to live for over a year following the craniotomy without exhibiting behavioral differences compared to non-operated controls, demonstrating the long-term stability of the procedure.

### Fluorescence imaging of neuronal activity

To capture calcium signals from resting-state neurons in mice, we utilized the RUSH system (Fan et al., 2019) to record the GCaMP6f fluorescence across the entire cranial window. The RUSH system comprises 35 sCMOS cameras arranged in a matrix to enable parallel acquisition of brain-wide calcium events. Briefly, this system is equipped with a custom objective lens, providing a field of view (FOV) of 1 × 1.2 cm² and a numerical aperture (NA) of 0.35. Images are captured by a camera array with a resolution of 14,000 pixels in height and 12,000 pixels in width. The overall magnification factor of the optical system is ×8. Fluorescence excitation occurs at 470 ± 20 nm, with emission detected at 525 ± 20 nm. In our experiments, we employed a frame rate of 4 fps. An external trigger signal synchronized the acquisition start time across cameras and different experimental stages.

### Intrinsic signal imaging for cortical mapping

Intrinsic signal imaging (ISI) measures the hemodynamic response of cortical tissue to visual stimuli across the entire field of view. This technique facilitates the generation of retinotopic maps that illustrate the spatial relationship between the visual field and the corresponding locations within each cortical area. These maps were used to delineate the boundaries of functionally defined visual areas, enabling precise analysis of the functional connectivity networks formed by neurons in the mouse visual cortex during the resting state (Figure 1a). The ISI methodology and data analysis procedures were based on the protocol outlined by Siegle et al. (2021).

### Experimental protocol and data acquisition

On the day of the experiment, the mouse’s head was secured in a custom-designed headframe, and the animal was allowed to acclimate in the dark for 10 minutes. Neural activity was then recorded from each experimental mouse for 30 minutes using the RUSH system. The first 10 minutes of recording were dedicated to capturing the mouse’s resting-state neural activity, during which no visual or other forms of stimuli were presented, and the mouse remained fully awake without engaging in any task. This period of data was used for analyzing the neural activity during the resting state. Subsequently, to facilitate neuronal localization, the mouse was exposed to 20 minutes of continuous visual stimulation, using a moving checkerboard pattern displayed on a 13-inch monitor. The monitor was positioned 15 cm from the mouse’s left eye, with the display rotated 30° clockwise along the mouse’s anterior-posterior axis and tilted 20° relative to the gravitational axis to ensure proper alignment of the left eye with the monitor’s plane.

The full 30-minute neural activity dataset was used for extracting neuronal data, while for the resting-state analysis, only the first 10 minutes of neural activity were used to construct the functional connectivity network. One week after fluorescence imaging, all experimental mice underwent ISI to precisely map the locations of the visual subareas. Prior to the experiment, all mice underwent at least 7 days of head fixation training to ensure they were in a relaxed and non-anxious state, minimizing potential stress-induced effects during the experiment. The data presented in this study were collected from a total of 8 mice (M1 to M8), which were used to investigate the neuronal network properties of the mouse primary visual cortex (V1) under resting-state conditions.

### Fluorescence image processing and neuron signal extraction

Given the high throughput of the RUSH system, we developed a customized parallel data analysis pipeline, adapted from CNMF-E (Zhou et al., 2018). Initially, the raw video data were motion-corrected through image registration. The corrected video was then temporally summarized into two key images: a pixel-neighbor correlation image and a peak-to-noise ratio image. The Hadamard product of these two images was used to identify initial neuronal candidates.

To refine the neuronal footprints and temporal activity, we applied constrained non-negative matrix factorization (CNMF) with a ring model to account for background noise. A vessel segmentation mask, generated based on intensity thresholds, was used to exclude neurons located on blood vessels. The temporal signals extracted from the neuronal candidates were then further processed using a supervised deep learning model, trained on manually curated data. This step effectively removed artifacts related to motion or hemodynamic fluctuations. Finally, after reliably extracting the neuronal signals, we aligned the neuronal footprints with the cranial window landmarks and mapped them to the visual areas identified through ISI. Using the Allen Mouse Brain Common Coordinate Framework version 3 (CCF v3) (Wang et al., 2020), we localized the neurons within specific cortical regions.

### Construction of functional connectivity network

In this study, we first performed band-pass filtering on the time-series data of mouse brain neurons to reduce the effects of low-frequency drift and high-frequency physiological noise. A band-pass frequency range of 0.1-1 Hz was selected to isolate the low-frequency signals associated with resting-state activity, while minimizing the influence of high-frequency interference and physiological noise.

After completing the data preprocessing, we constructed a functional brain network based on the correlation between neuronal time series. A correlation matrix was generated by calculating the Pearson correlation coefficient for each pair of neuronal time series. Each element of the matrix represented the linear relationship between a pair of neurons, with values ranging from −1 (perfect negative correlation) to 1 (perfect positive correlation). To enhance the tractability of the network structure, we first took the absolute value of the correlation matrix and then applied a threshold to construct a binary matrix. This approach ensured that connections with greater strength, whether positively or negatively correlated, were included in the network analysis. We selected a threshold range from 0 to 0.4 to systematically explore the influence of different threshold levels on the topological properties of the functional connectivity networks. At each threshold level, connections in the correlation matrix were retained if their absolute values exceeded the set threshold, ensuring that the constructed network preserved critical connections while minimizing over-connections.

### Analysis of network properties analysis

After constructing the functional connectivity network, we performed a comprehensive analysis that included small-world network analysis, rich-club organization analysis, and modularity maximization analysis. Constrained by computational resources, we performed the calculations using 10 randomly generated networks (van den Heuvel et al., 2008).

**Analysis of small-world network properties.** Four small-world parameters (clustering coefficient *C*, characteristic path length *L*, normalized clustering coefficient γ, and normalized shortest path length λ) were adopted to characterize the global topological organization of brain networks. For a given graph G with N nodes, the clustering coefficient *C* is defined as (Tian et al., 2011; Watts & Strogatz, 1998):

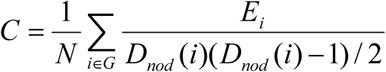

Here *D*_*nod*_ (*i*) denotes the degree of node *i*, defined as the number of edges connected to that node, and *E_i_* represents the total number of edges within the subgraph formed by the neighbors of node *i*. The characteristic path length *L* is defined as follows (Newman, 2003; Tian et al., 2011):

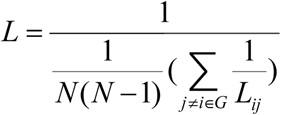

Of these two parameters, the clustering coefficient *C* quantifies the local interconnectedness within a network, while the characteristic path length *L* reflects the overall efficiency of information routing across the network.

To investigate the small-world properties, the normalized clustering coefficient *γ =C_net_/C_random_* and normalized characteristic path length *λ =L_net_/L_random_* were calculated (Watts & Strogatz, 1998). In these equations, *C_net_* and *L_net_* represent the clustering coefficient and characteristic path length of the real network, respectively,, while *C_random_* and *L_random_* denote the corresponding values for the degree-matched random networks, which preserve the same number of nodes, edges, and degree distribution (Maslov & Sneppen, 2002). A network is considered to exhibit small-world properties if it satisfies the conditions γ > 1 and λ ≈ 1 (Watts & Strogatz, 1998), or equivalently, if the small-world index σ = γ / λ > 1 (Humphries et al., 2006).

**Examination of rich-club organization.** For a given graph G, the degree of each node *i* in the network was determined by counting the number of edges connecting node *i* to other nodes. A threshold degree *k* was defined, and nodes with a degree less than or equal to *k* were excluded from the network. The degree *k* represents the minimum number of connections a node must have to remain in the network. The rich-club coefficient *ф(k)* for the remaining network was then calculated as the ratio of the actual number of connections between the remaining nodes to the total possible number of connections if all remaining nodes were fully interconnected. Formally, the rich-club coefficient *ф(k)* is defined as follows (Colizza et al., 2006; McAuley et al., 2007; Zhou & Mondragón, 2004):

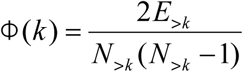

The rich-club coefficient ф(k) quantifies the extent to which nodes with a degree greater than a given threshold *k* are more densely interconnected than would be expected in a random network. Specifically, *N* > *_k_* denotes the number of nodes with degree strictly greater than k, and *E* > *_k_* represents the actual number of connections (edges) among these high-degree nodes. The denominator, *N* > *_k_* (*N* > *_K_* –1) / 2, corresponds to the maximum possible number of edges that could exist among the *N* > *_k_* nodes if they formed a fully connected subgraph. By normalizing the actual edge count using this theoretical maximum, *ф(k)* provides a relative measure of interconnectivity among the high-degree nodes. A higher value of *ф(k)* indicates a stronger “rich-club” organization, wherein high-degree nodes preferentially form tightly interconnected communities.

**Modularity detection and maximization.** The community structure of the network was identified using the Louvain algorithm (Blondel et al., 2008), a widely adopted method for detecting modularity by maximizing within-community connections relative to a null model. After partitioning the network into distinct communities, the quality of the detected modular organization was evaluated by computing the modularity index Q, defined as (Sporns & Betzel, 2016):

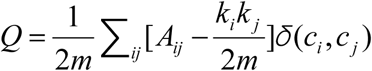

In this equation, *A_ij_* represents the weight of the connection (edge) between nodes *i* and *j,* while *k_i_* and *k _j_* denote the degrees of nodes *i* and *j,* respectively. The variable *m* indicates the total number of edges present in the network. The function *δ*(*c_i_*, *c _j_*) is an indicator function that equals 1 if nodes *i* and *j* belong to the same community and equals 0 otherwise. The term 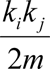 represents the expected number of edges between nodes *i* and *j* under a null model, where edges are randomly assigned while preserving the degree distribution of the original network. This null model provides a baseline for assessing whether observed within-community connections are significantly greater than expected by chance. Therefore, the modularity index Q quantifies the difference between the actual density of edges within communities and the expected edge density derived from the random model. Higher values of Q indicate a more prominent and clearly delineated community structure, typically ranging from 0 (absence of community structure) to 1 (highly segregated communities).

## Supplemental Material

**Supplementary Figure 1.**
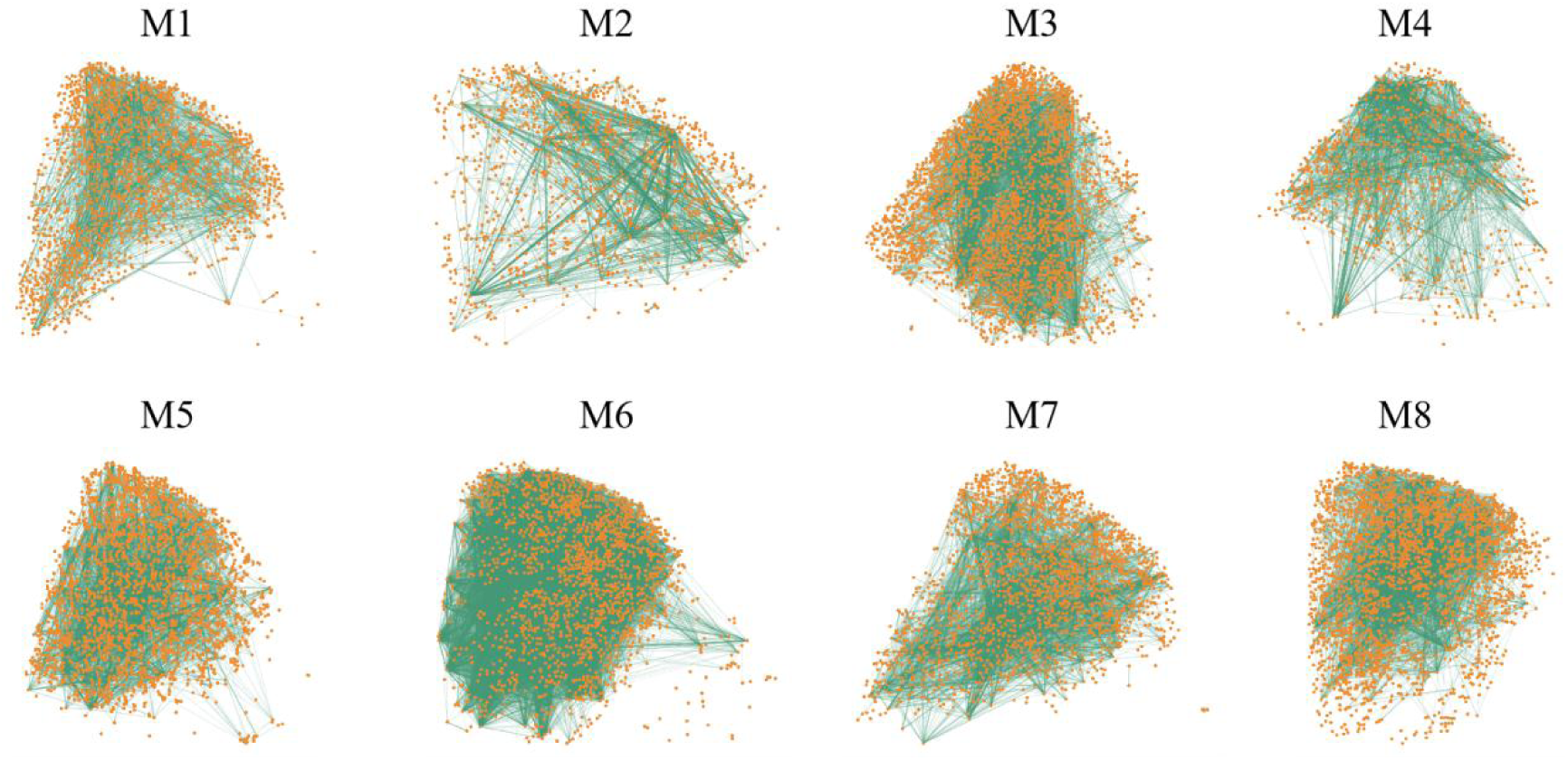
Functional connectivity networks from eight representative mice constructed at threshold T = 0.50. This figure shows the neuronal network architecture based on spontaneous activity recorded from approximately 2,000 neurons in the mouse primary visual cortex (V1) during the resting state. Each panel (M1 to M8) corresponds to one of the eight representative mice in the study. The orange dots represent individual neurons, while the green lines represent functional connections between neurons based on Pearson correlation coefficients. As the correlation threshold increases (T = 0.50), weaker connections are pruned, but the core structure of the network, including dense clustering and hub-like neurons, remains intact. These observations highlight the stable organization and modularity of the V1 network, which reflects efficient integration across distinct functional sub-networks in the cortical microcircuit. The maintenance of structural features across varying mice emphasizes the robustness of these network properties at the mesoscopic scale.

**Supplementary Figure 2.**
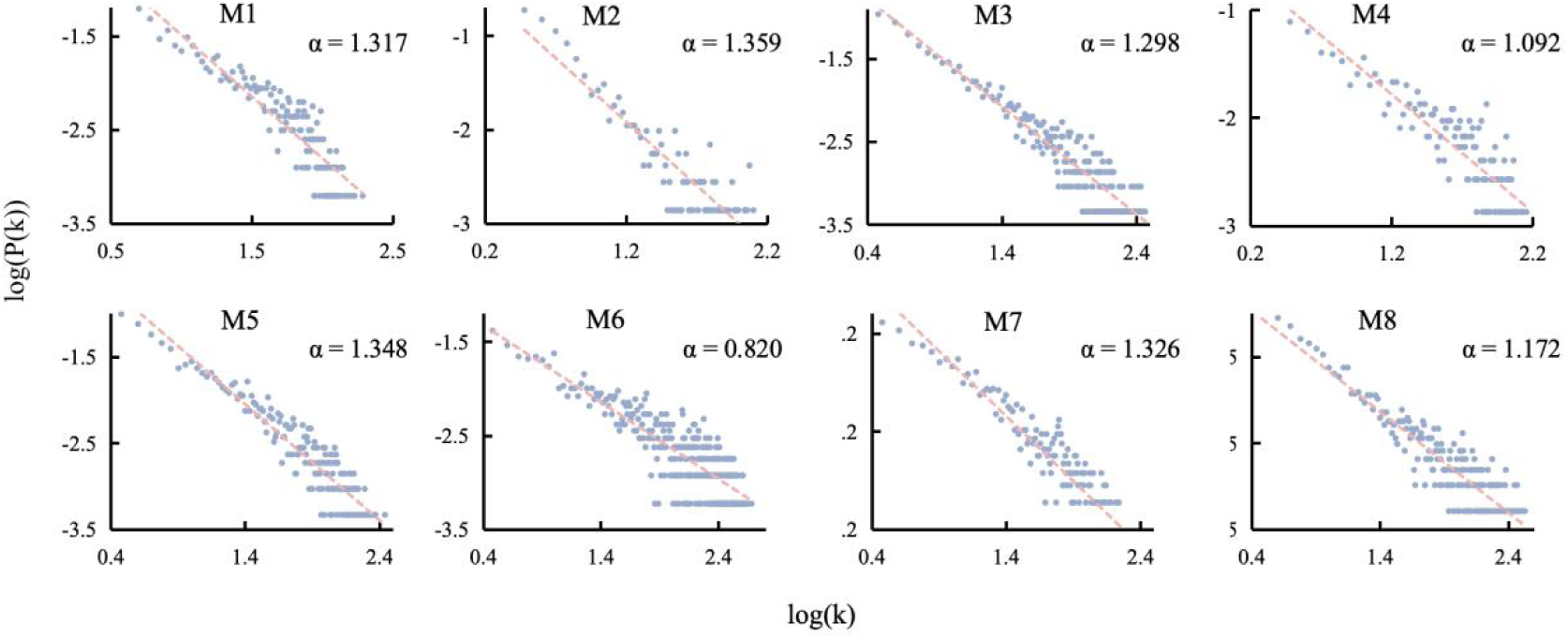
Degree distributions of functional connectivity networks from eight mice. This figure shows the log-log plots of the degree distributions of neuronal networks at a correlation threshold of T = 0.40 for eight individual mice (M1 to M8). The x-axis represents the logarithmic values of node degrees (k), while the y-axis shows the logarithmic probability distribution of the node degrees (P(k)). Each panel displays the degree distribution for one mouse, with the red dashed lines representing the best-fit power-law distributions. The power-law exponents (α) for each distribution are provided within each panel. These distributions exhibit the characteristic heavy-tailed, scale-free nature, where a small subset of neurons (hubs) possess significantly higher degrees compared to the majority of the network. The fitted power-law exponents for the degree distributions range from α = 0.820 (M6) to α = 1.359 (M2), confirming the presence of scale-free network properties across different mice. These results indicate that the V1 neuronal networks follow a hub-centric organization, with a small number of highly connected neurons playing a crucial role in network integration.

**Supplementary Figure 3.**
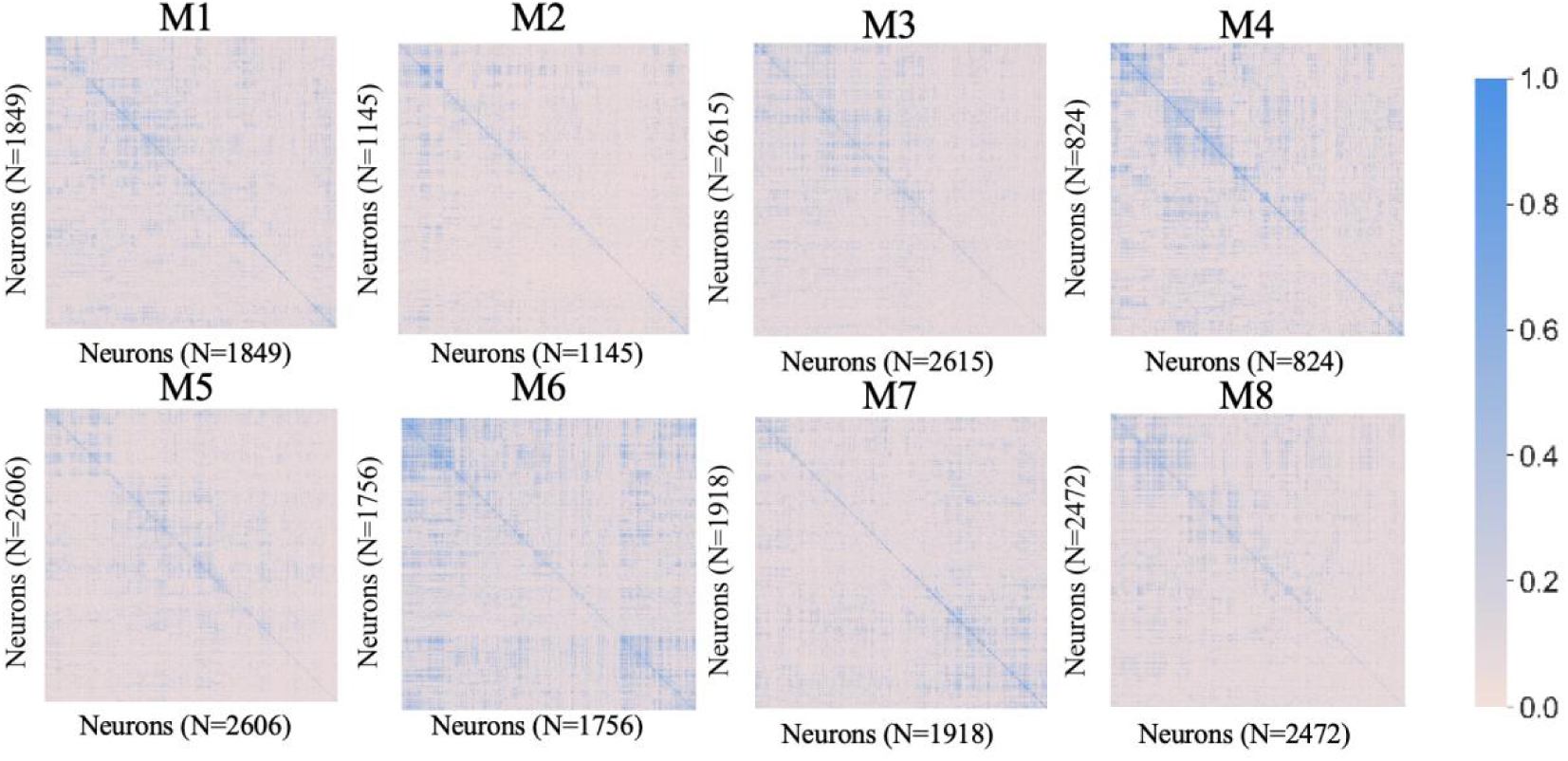
Functional connectivity matrices of eight mice. This figure presents the pairwise Pearson correlation matrices for neuronal activity from eight individual mice (M1 to M8). Each matrix shows the strength of functional connectivity between all pairs of neurons within the primary visual cortex (V1) during the resting state, with the color scale indicating the correlation values (ranging from 0 to 1, with darker blue indicating stronger correlations). The number of neurons recorded from each mouse is indicated in parentheses (N = 824 to N = 2615). The dense diagonal structure in each matrix reflects the self-correlation of neurons, while off-diagonal elements represent the correlations between different neuronal pairs. The presence of distinct patterns of connectivity in each matrix highlights the modular organization of the V1 network, with neurons clustering into regions of high correlation. These results provide further evidence for the modular and organized structure of neuronal networks in V1, which supports the hypothesis of a highly structured functional connectivity architecture within the cortical microcircuit.

## Acknowledgments

We sincerely thank Xiaojuan Wu for her assistance in animal preparation, and Yun Chen for his support during data acquisition. We also appreciate the valuable suggestions provided by Tao Liu during the preparation of the figures.

