## Supplemental Figure for "Extreme Small-World, Modular, and Rich-Club Topology of Single-Neuron Networks in Mouse Primary Visual Cortex"

**Supplemental Material**

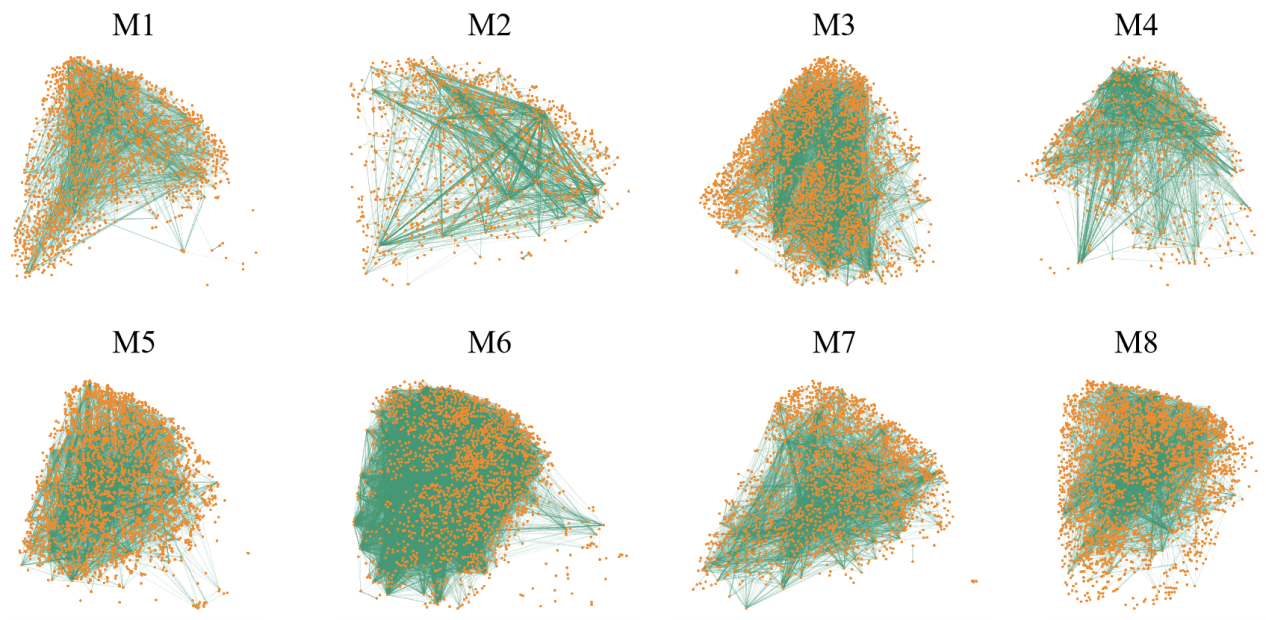

**Supplementary Figure 1.** **Functional connectivity networks from eight representative mice constructed at threshold T = 0.50.** This figure shows the neuronal network architecture based on spontaneous activity recorded from approximately 2,000 neurons in the mouse primary visual cortex (V1) during the resting state. Each panel (M1 to M8) corresponds to one of the eight representative mice in the study. The orange dots represent individual neurons, while the green lines represent functional connections between neurons based on Pearson correlation coefficients. As the correlation threshold increases (T = 0.50), weaker connections are pruned, but the core structure of the network, including dense clustering and hub-like neurons, remains intact. These observations highlight the stable organization and modularity of the V1 network, which reflects efficient integration across distinct functional sub-networks in the cortical microcircuit. The maintenance of structural features across varying mice emphasizes the robustness of these network properties at the mesoscopic scale.

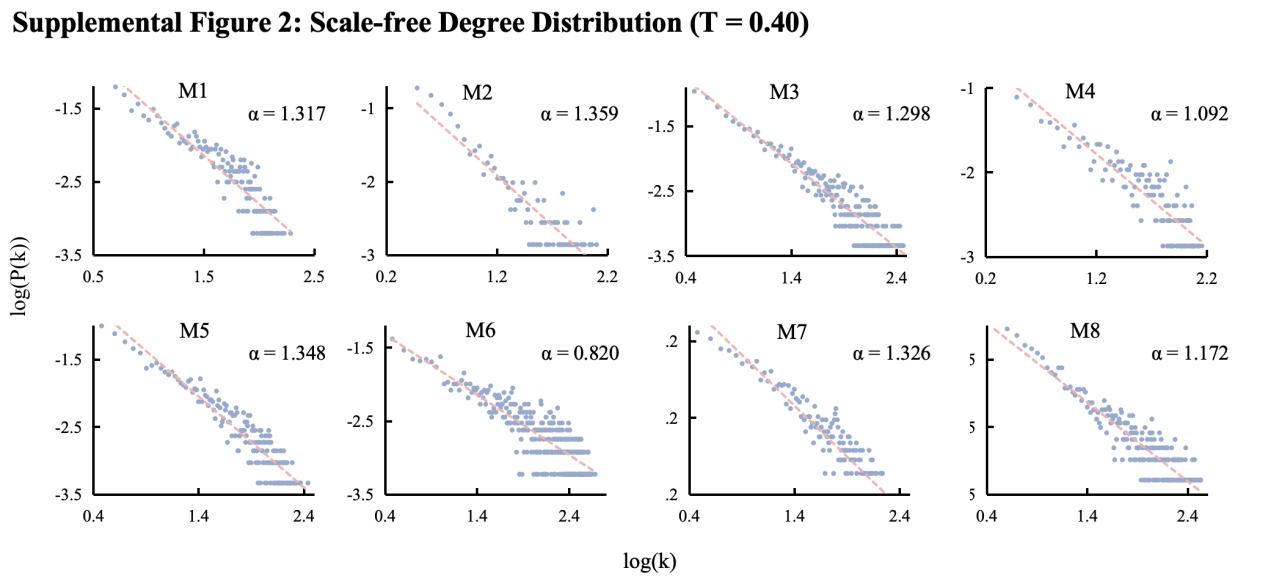

**Supplementary Figure 2.** **Degree distributions of functional connectivity networks from eight mice.** This figure shows the log-log plots of the degree distributions of neuronal networks at a correlation threshold of T = 0.40 for eight individual mice (M1 to M8). The x-axis represents the logarithmic values of node degrees (k), while the y-axis shows the logarithmic probability distribution of the node degrees (P(k)). Each panel displays the degree distribution for one mouse, with the red dashed lines representing the best-fit power-law distributions. The power-law exponents (α) for each distribution are provided within each panel. These distributions exhibit the characteristic heavy-tailed, scale-free nature, where a small subset of neurons (hubs) possess significantly higher degrees compared to the majority of the network. The fitted power-law exponents for the degree distributions range from α = 0.820 (M6) to α = 1.359 (M2), confirming the presence of scale-free network properties across different mice. These results indicate that the V1 neuronal networks follow a hub-centric organization, with a small number of highly connected neurons playing a crucial role in network integration.

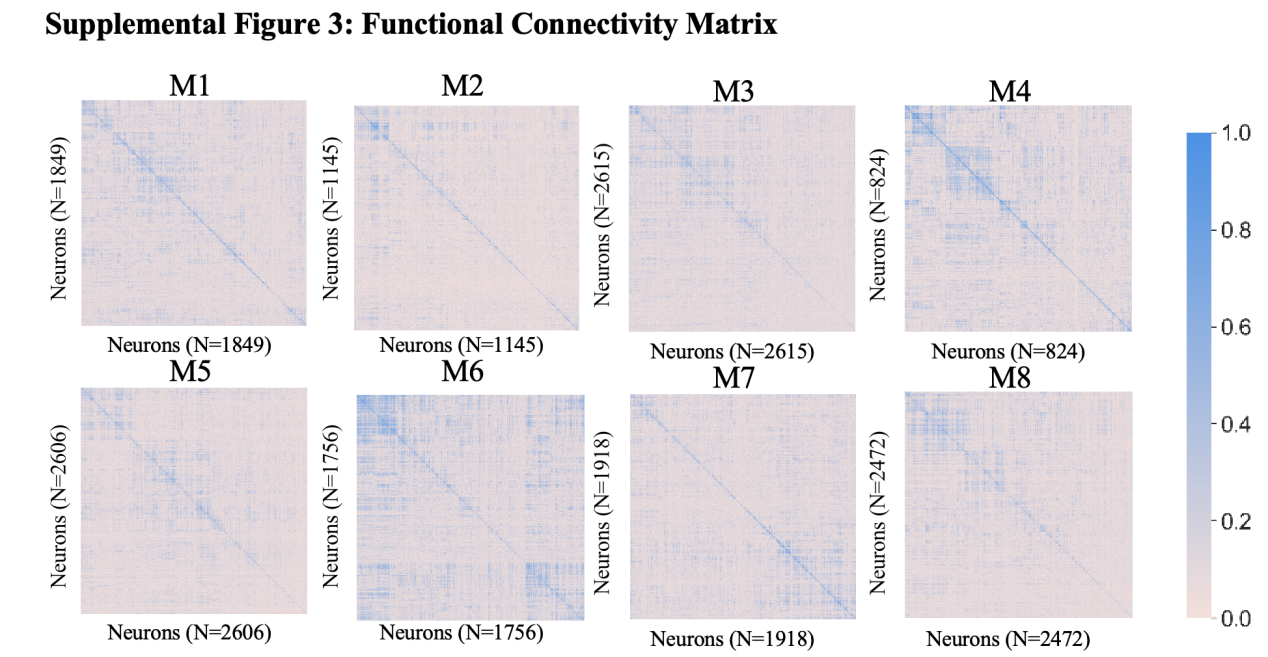

**Supplementary Figure 3. Functional connectivity matrices of eight mice.** This figure presents the pairwise Pearson correlation matrices for neuronal activity from eight individual mice (M1 to M8). Each matrix shows the strength of functional connectivity between all pairs of neurons within the primary visual cortex (V1) during the resting state, with the color scale indicating the correlation values (ranging from 0 to 1, with darker blue indicating stronger correlations). The number of neurons recorded from each mouse is indicated in parentheses (N = 824 to N = 2615). The dense diagonal structure in each matrix reflects the self-correlation of neurons, while off-diagonal elements represent the correlations between different neuronal pairs. The presence of distinct patterns of connectivity in each matrix highlights the modular organization of the V1 network, with neurons clustering into regions of high correlation. These results provide further evidence for the modular and organized structure of neuronal networks in V1, which supports the hypothesis of a highly structured functional connectivity architecture within the cortical microcircuit.
